# Analysis and correction of compositional bias in sparse sequencing count data

**DOI:** 10.1101/142851

**Authors:** M. Senthil Kumar, Eric V. Slud, Kwame Okrah, Stephanie C. Hicks, Sridhar Hannenhalli, Héctor Corrada Bravo

## Abstract

Count data derived from high-throughput DNA sequencing is frequently used in quantitative molecular assays. Due to properties inherent to the sequencing process, unnormalized count data is compositional, measuring *relative* and not *absolute* abundances of the assayed features. This *compositional bias* confounds inference of absolute abundances. We demonstrate that existing techniques for estimating compositional bias fail with sparse metagenomic 16S count data and propose an empirical Bayes normalization approach to overcome this problem. In addition, we clarify the assumptions underlying frequently used scaling normalization methods in light of compositional bias, including scaling methods that were not designed directly to address it.

## Background

Sequencing technology has played a fundamental role in 21st century biology: the output data, in the form of sequencing reads of molecular features in a sample, are relatively inexpensive to produce [1, 2, 3, 4]. This, along with the immediate availability of effective, open source computational toolkits for downstream analysis [5, 6], has enabled biologists to utilize this technology in ingenious ways to probe various aspects of biological mechanisms and organization ranging from microscopic DNA binding events [7, 8] to large-scale oceanic microbial ecosystems [9, 10].

This remarkable flexibility of sequencing comes with atleast one tradeoff. As noted previously in the literature [11, 12, 13, 14] (illustrated in **Fig. 1**), unnormalized counts obtained from a sequencer only reflect relative abundances of the features in a sample, and not their absolute internal concentrations. When a differential abundance analysis is performed on this data, fold changes of null features, those not differentially abundant in the *absolute* scale, are intimately tied to those of features that are perturbed in their absolute abundances, making the former appear differentially abundant. We refer to this artifact as *compositional bias*. Such effects are observable in the count data from the large-scale Tara oceans metagenomics project [10], (**Fig. 2**), in which a few dominant taxa are attributable to global differences in the between-oceans fold-change distributions.

**Figure 1.**
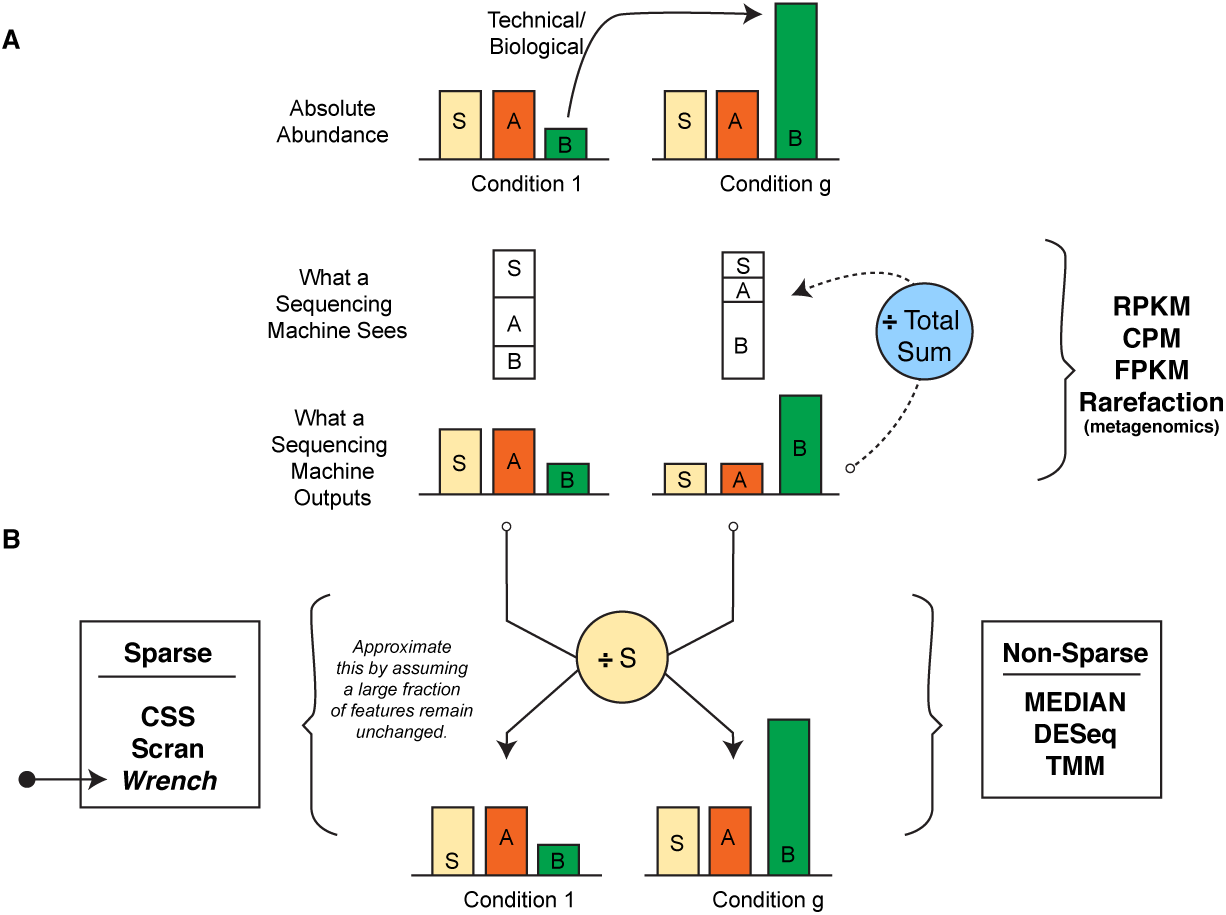
Scaling normalization approaches from the perspective of compositional correction. (**A**) Features *S* and *A* have similar absolute abundances in two experimental conditions, while *B* has increased in its absolute abundance in condition *g* due to technical/biological reasons. Because of the proportional nature of sequencing, increase in *B* leads to reduced read generation from others (compositional bias). An analyst would reason *A* and *S* to be significantly reduced in abundance, while, in reality they did not. (**B**) Knowing *S* is expressed at the same concentration in both conditions allows us to scale by its abundance, resolving the problem. DESeq and TMM, by exploiting rerefence strategies across feature count data (described below), approximate such a procedure, while techniques that are based only on library size alone like RPKM and rarefication/subsampling can lead to unbiased inference only under very restrictive conditions. Approaches available for sparse settings are indicated. *Wrench* is the proposed technique in this paper.

**Figure 2.**
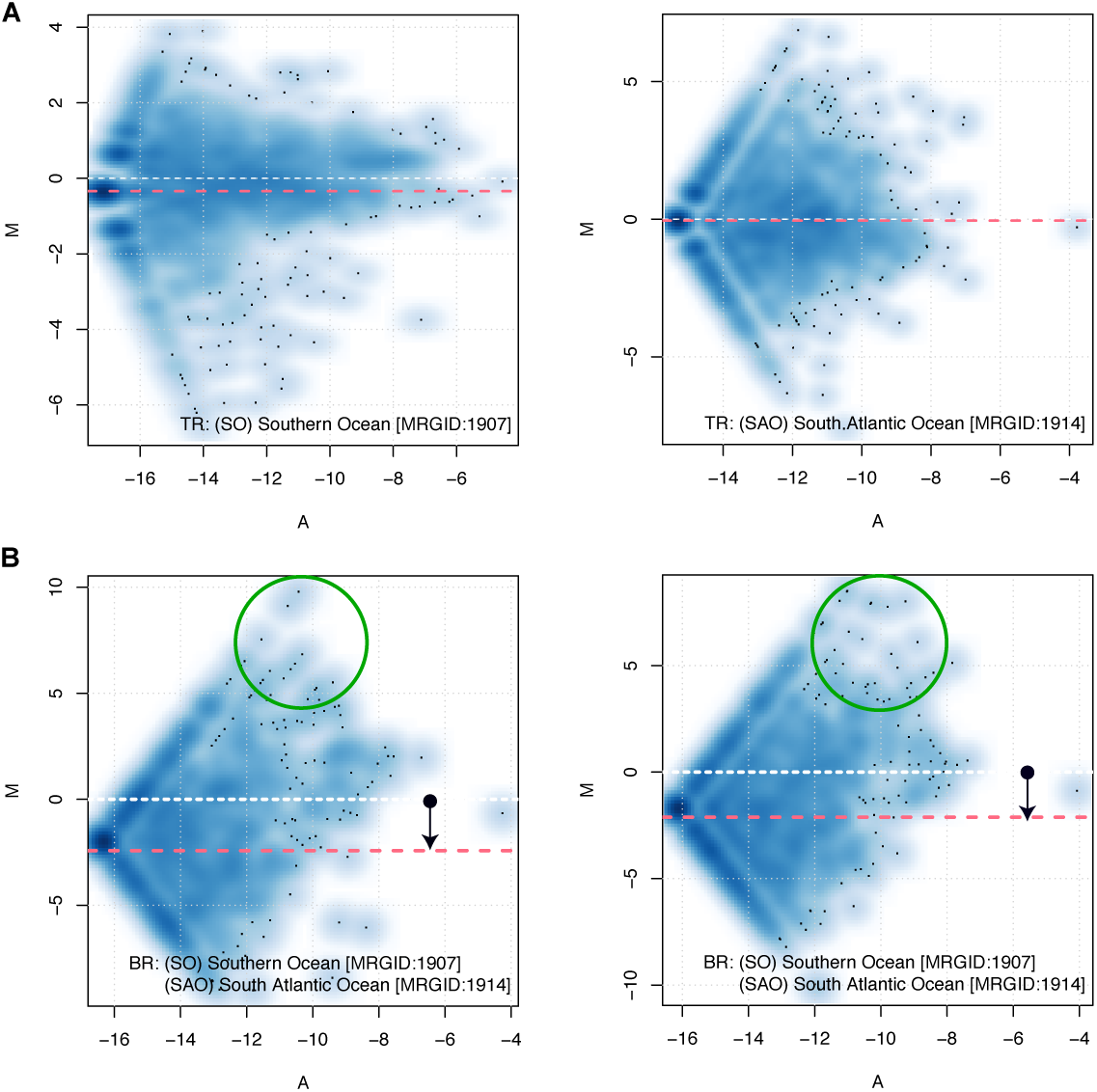
Importance of compositional bias correction in sparse metagenomic data. (A) M-A pots of 16S reconstructions (from high sequencing depth, whole metagenome shotgun sequencing experiments) from two technical replicates each from the Tara oceans project [10] generated for the Southern and South Atlantic Oceans. In all subplots, x-axis plots for each feature, its average of the logged proportions in the two compared samples; y-axis plots the corresponding differences. The red dashed line indicates the median log fold change, which is 0 across the technical replicates. (B) M-A plots of the same replicates but plotted across the two oceans. The median of the log-fold change distribution is clearly shifted. A few dominant taxa in the South Atlantic Ocean (circled) are attributable for driving this overall apparent differences in the observed fold changes. The Tara 16s dataset, reconstructed from very deep whole metagenome shotgun experiments of oceanic samples, albeit boasting of an average 100,000 16S contributing reads per sample, still encourages a median 88% feature absence per sample.

Correction for compositional bias can be achieved by re-scaling each sample’s count data with its corresponding count of an internal control feature (or “spike-in”, **Fig. 1B**). In the absence of such control features, effective correction for compositional bias can still be hoped for, as it can be shown that this correction amounts to resolving a linear technical bias [13]. This fact allows one to exploit several widely used non-spike-in normalization approaches [15, 13, 16, 17], which approximate the aforementioned spike-in strategy by assuming that most features do not change on average across samples/conditions. For the same reason, such an interpretation can also be given to approaches like centered logarithmic transforms (CLR) from the theory of compositional data, which many analysts favor when working with relative abundances [18, 19, 20, 21, 22, 23, 24]. In this paper, we analyze the behavior of these existing scaling normalization techniques in light of compositional bias.

When trying to normalize metagenomic 16S survey data with these methods however, we found that the large fraction of zeroes in the count data, and the relatively low sequencing depths of metagenomic samples posed a severe problem: DESeq failed to provide a solution for all the samples in a dataset of our interest, and TMM based its estimation of scale factors on very few features per sample (as low as 1). The median approach simply returned zero values. CLR transforms behaved similarly. When one proceeds to avoid this problem by adding pseudo-counts, owing to heavy sparsity underlying these datasets, the transformations these techniques imposed mostly reflected the value of pseudocount and the number of features observed in a sample. A recently established scaling normalization technique, Scran [25], tried to overcome this sparsity issue in the context of single cell RNAseq count data – which also entertains a large fraction of zeroes – by decomposing simulated pooled counts from multiple samples. That approach, developed for relatively high coverage single cell RNAseq, also failed to provide solutions for a significant fraction of samples in our datasets (as high as 74%). Furthermore, as we illustrate later, compositional bias affects data sparsity, and normalization techniques that ignore zeroes when estimating normalization scales (like CSS [26], and TMM) can be severely biased. The relatively low sequencing depth per sample (as low as 2000 reads per sample), large number of features and their diversity across samples thus pose a serious challenge to existing normalization techniques. In this paper, we develop a compositional bias correction technique for sparse count data based on an empirical Bayes approach that borrows information across features and samples

Since we have presented the problem of compositional bias as one affecting inferences on absolute abundances, one might wonder if resolving compositional bias is needed when analyses on *relative* abundances are performed. It is important to realize that compositional bias is infused in the count data, solely due to inherent characteristics of the sequencing process, even before it passes through any specific normalization process like scaling by library size. In practical conditions, because feature-wise abundance perturbations are also driven by technical sources of variation uncorrelated with total library size [27, 28, 29, 30], compositional bias correction becomes necessary even when analysis is performed on relative abundances.

The paper is organized as follows. We first set up the problem of compositional bias correction and with appropriate simulations, evaluate several scaling normalization techniques in solving it. We find that techniques based only on library size (e.g. unaltered RPKM/CPM [31], rarefication/subsampling in metagenomics [32, 33]) are provably bad. Other scaling techniques, while providing robust compositional bias estimates on high coverage data, perform poorly at sparsity levels often observed with metagenomic count data. We then introduce the proposed normalization approach (*Wrench*) and evaluate its performance with simulations and experimental data showing that it can lead to reduced false positives and rich annotation discoveries. We close by discussing the insights obtained by applying Wrench and other scaling normalization techniques to experimental datasets, arguing both for addressing compositional bias in general practice and in benchmarking studies. Because all the aforementioned techniques, including our own proposal, assume that most features do not change across conditions on average, they would all suffer in analyses of features arising from arbitrary general conditions. In such cases, spike-in based techniques can be effective [34], although methods similar to the ERCC method for bulk RNAseq will not work for the simple reason it starts with an extract, an already compositional data source.

## Results

### Formalizing compositional bias in differential abundance analysis

Below, we describe the *compositional correction factor*, the quantity we use to evaluate scaling normalization techniques in overcoming compositional bias.

**Fig. 3** illustrates a general sequencing experiment and sets up the problem of compositional bias correction. We imagine a set of samples/observations *j* = 1 … *n*_*g*_ arising from conditions *g* = 1 … *G* (e.g., cases and controls). The true absolute abundances of features in every sample organized as a vector 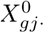, are perturbed by various technical sources of variation as the sample is prepared for sequencing. The end result is a transformed absolute abundance vector *X*_*gj*_., the net total abundance of which is denoted by *T*_*gj*_ = ∑_*i*_ *X*_*gji*_ = *X*_*gj*+_, where the + indicates summing over that subscript. This is the input to the sequencer, which introduces compositional bias by producing reads *proportional* to the *absolute* feature abundances represented in *X*_*gj*_‥ The output reads are processed and organized as counts in a vector *Y*_*gj*_., which now retain only *relative* abundance information of features in *X*_*gj*_‥ The ultimate goal of a normalization strategy is to recover 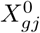. for all *g* and *j*.

**Figure 3.**
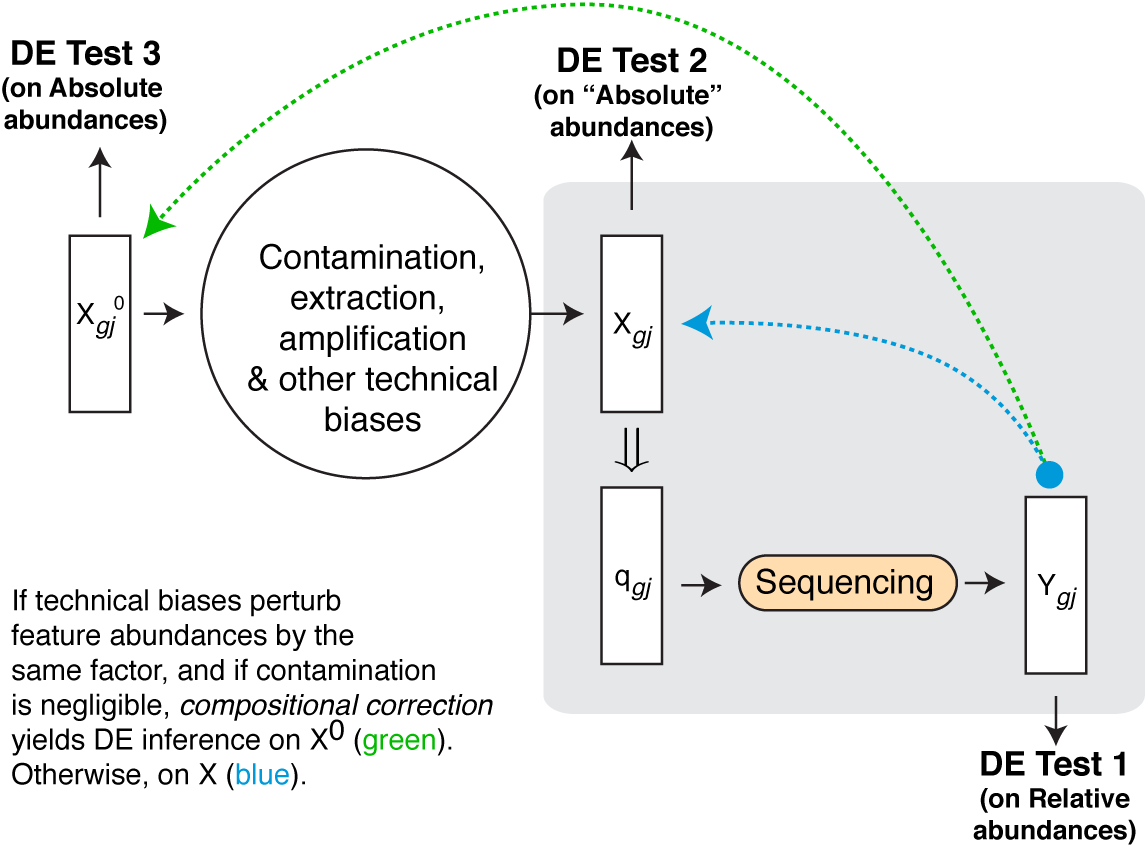
Compositional bias introduced by sequencing technology. As a sample *j* from group *g* of interest is prepared for sequencing, its true internal feature concentrations (organized as a vector) 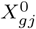 is transformed by various technical biases to *X*_*gj*_. A sequencing machine introduces compositional bias by generating counts *Y*_*gj*_ proportional to the input absolute abundances in *X*_*gj*_ according to proportions *q*_*gj*_ = [… *x*_*gji*_/(∑_*k*_ *x*_*gjk*_)…], *i* and *k* indexing features. Directly performing a differential abundance test on *Y* (DE Test 1), by using normalization factors proportional to that of total sequencing output (ex: R/FPKM/subsampling in metagenomics) amounts to testing for changes in *relative* abundances of features in *X*, in general (*not X*^0^). For inferring differences in absolute abundance, we need to reconstruct *X*^0^ from *Y* to perform our inference (DE Test 3). For compositional bias correction in particular, we care about reconstructing *X*_*j*_ from *Y* (DE Test 2). We show more formally later that compositional correction can reconstruct *X*^0^ if technical biases perturb all feature abundances by the same factor, and that the presence of sequence-able contaminants induces more stricter assumptions behind their application.

Our goal is to evaluate existing normalization approaches based on how well they reconstruct *X* from *Y*, as it is in this step, that the sequencing process induces the bias we are interested in. We come back to the question of reconstructing *X*^0^ at the end of this subsection. Because we are ignoring all other technical biases inherent to the experiment/technology (i.e., the process from *X*^0^ → *X*), our discussions apply to RNAseq/scRNAseq/metagenomics and other quantititative sequencing based assays. In this paper, our primary interest will be in the correction of compositional bias for metagenomic marker gene survey data, which are often under-sampled.

Although not strictly necessary, for simplicity, we shall assume that the relative abundances of each feature *i* is given by *q*_*gi*_ for all samples within a group *g*. It is also reasonable to assume an *X*_*gj*_.|*T*_*gj*_ ~ *Multinomial*(*T*_*gj*_,*q*_*g*_.), where *q*_*g*_. is the vector of feature-wise relative abundances. Such an assumption follows for example from a Poisson assumption on the expression of features *X*_*gji*_ [35]. Similarly, we shall assume the observed counts 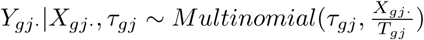, *τ*_*gj*_ is the corresponding sampling depth. Notice that marginally, *E*[*Y*_*gji*_|*τ*_*gj*_] = *q*_*gi*_ · *τ*_*gj*_, and hence averaging the observed sample-wise proportions 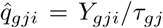 in group *g* for feature *i* yields the marginal expectation 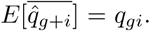. We shall use *E*[*T*_*g*1_] to denote the average (across samples) *total* absolute abundance of features in group *g* at the time of input. Similarly, 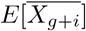 will denote the marginal expectation of absolute abundance of feature *i* across samples in group *g* (number of molecules per unit volume in case of RNAseq / number of distinct 16S fragments per unit volume in an environmental lysate in the case of 16S metagenomics). If we set *g* = 1 as the control group, and define, for every feature *i*, 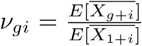, then log*v*_*gi*_ is the log-fold change of true absolute abundances associated with group *g* relative to that of the control group. We can write:

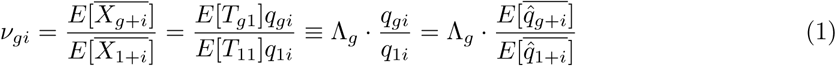

This indicates that the fold changes based on observed proportions (estimated from *Y*) from the sequencing machine confounds our inference of the fold changes associated with absolute abundances of features at stage *X*, through a linear bias term Λ_*g*_. Thus, to reconstruct the average absolute abundances of features in experimental group *g*, one needs to estimate the *compositional correction factor* 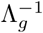, where for convenience in exposition below, we have chosen to work with the inverse. Note that the compositional correction factor for the control group 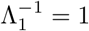 by definition.

Details on our terminology and how it differs from *normalization factors*, which are compositional factors altered by sample depths, are presented in the Simulations subsection under Methods. Below, we use the terms compositional scale or more simply scale factor interchangeably to refer to compositional correction factors.

*The central idea in estimating compositional correction factors* For any group *g*, an effective strategy for estimating 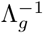 can be derived based on an often quoted assumption behind scale normalization techniques [13]: if most features do not change in an experimental condition relative to the control group, eqn. 1 should hold true for most features with *v*_*gi*_ = 1. Thus, an appropriate summary statistic of these ratios of proportions could serve as an estimate of 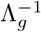.

So far we have discussed estimating group-specific compositional factors. With this idea in place, a normalization procedure for deriving sample-specific compositional scale factors can be devised. One only needs to carry out the above procedure by pretending that every sample arises from its own experimental group. Indeed, as illustrated in **Table 1**, many scale normalization methods (including the proposal in this work) can be viewed in this light, where some control set of proportions (“reference”) is defined, and the 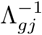 estimate is derived for every sample *j* based on the ratio of its proportions to that of the reference. This central idea being the same, the robustness of these methods are dependent on how well the assumptions hold with respect to the chosen reference, and the choice of the estimation strategy.

**Table 1.** Scaling normalization approaches derive their technical bias estimates from ratio of proportions. For each scaling normalization technique (rows of the table, named in the first column), we present the transformation they apply to the raw count data (second column) to produce normalize counts. The third column shows how all techniques use statistics based on ratio of proportions (third column) to derive their scale factors. In the table, *i* = 1… *p* indexes features (genes/taxonomic units), and each sample is considered to arise from its own singleton group: *g* = 1 … *n* and *j* = 1, *τ*_*gj*_ the sample depth of sample *j*, *q*_*gji*_ the proportion of feature *i* in sample *j*, *w*_*ij*_ represents a weight specific to each technique, and 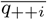 is the average proportion of feature *i* across the dataset. In the second column, the first row in each cell represents the transformation applied on the raw count data by the respective normalization approach. They all adjust a sample’s counts based on sample depth (*τ*_*gj*_) and a compositional scale factor 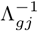. As noted in the third column, the estimation of 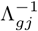 is based on the ratio of sample-wise relative abundances/proportions (*q*_*gji*_) to a reference that are all some robust measures of central tendency in the count data. The logarithmic transform accompanying CLR should not worry the reader about its relevance here, in the following sense: the log-transformation often makes it possible to apply statistical tests based on normal distributions for the rescaled data; this is in-line with applying log-normal assumptions on the rescaled data obtained with the rest of the techniques. *C* = [∏_*j*_ *τ*_*gj*_]^−1/*n*^ is a constant factor independent of sample, and its presence does not matter. For the same reason, Median and Upper Quartile scalings and CLR transforms, can be thought to base their estimates on a reference that assigns equal mass to all the features or if the reader wishes, a more complicated reference that behaves proportionally. When most features are zero, values arising from classical scale factors can be severely biased or undefined as we shall illustrate in the rest of the paper.

| Technique | Proposed Abundance Measure, Scale factor | Signal for Compositional Scale in |
| --- | --- | --- |
| Total Sum | $\frac{y_{gji}}{\tau_{gj} \cdot \Lambda_{gj}^{-1}},$<br>$\Lambda_{gj}^{-1} = 1$ | |
| TMM | $\frac{y_{gji}}{\tau_{gj} \cdot \Lambda_{gj}^{-1}},$<br>$\Lambda_{gj}^{-1} = e^{\left[ \sum_{i: y_{ij} > 0} \cap i \in \text{trimmed set for } j} w_{ij} \log \left( \frac{q_{gji}}{q_{1ji}} \right) \right]}$ | $\frac{q_{gji}}{q_{1ji}},$ ratio of proportions |
| DESeq | $\frac{y_{gji}}{C \cdot \tau_{gj} \cdot \Lambda_{gj}^{-1}} \propto \frac{y_{gji}}{\tau_{gj} \cdot \Lambda_{gj}^{-1}},$<br>$\Lambda_{gj}^{-1} = \text{median}_i \frac{q_{gji}}{[\prod_k q_{ik}]^{\frac{1}{n}}}$ | $\frac{q_{gji}}{[\prod_k q_{ik}]^{\frac{1}{n}}},$ ratio of proportions |
| Median | $\frac{y_{gji}}{\tau_{gj} \cdot \Lambda_{gj}^{-1}},$<br>$\Lambda_{gj}^{-1} = \text{median}_i q_{gji} \propto \text{median}_i \frac{q_{gji}}{1/p}$ | $\frac{q_{gji}}{1/p},$ ratio of proportions |
| Upper quartile | $\frac{y_{gji}}{\tau_{gj} \cdot \Lambda_{gj}^{-1}},$<br>$\Lambda_{gj}^{-1} = \text{upper quartile}_i q_{gji} \propto \text{upper quartile}_i \frac{q_{gji}}{1/p}$ | $\frac{q_{gji}}{1/p},$ ratio of proportions |
| CLR Transformation | $\log \left( \frac{y_{gji}}{[\prod_i y_{gji}]^{\frac{1}{p}}} \right) \equiv \log \left( \frac{q_{gji}}{[\prod_i q_{gji}]^{\frac{1}{p}}} \right) \equiv \log \left( \frac{y_{gji}}{\tau_{gj} \cdot \Lambda_{gj}^{-1}} \right),$<br>with $\Lambda_{gj}^{-1} = [\prod_i q_{gji}]^{\frac{1}{p}} \propto [\prod_i \frac{q_{gji}}{1/p}]^{\frac{1}{p}}$ | $\frac{q_{gji}}{1/p},$<br>closely tracks Median factors above;<br>ratio of proportions |
| Scran | $\frac{y_{gji}}{\tau_{gj} \cdot \Lambda_{gj}^{-1}},$<br>$\Lambda_{gj}^{-1} = \text{fit linear models to } \left\{ \frac{q_{1ji}}{q_{++i}}, \dots, \frac{q_{nji}}{q_{++i}} \right\}_{i=1}^p$ | $\frac{q_{gji}}{q_{++i}},$ ratio of proportions |
| Wrench | $\frac{y_{gji}}{\tau_{gj} \cdot \Lambda_{gj}^{-1}},$<br>$\Lambda_{gj}^{-1} = \frac{1}{p} \sum_i w_{ij} \frac{q_{gji}}{q_{++i}}$ | $\frac{q_{gji}}{q_{++i}},$ ratio of proportions |

*Reconstrucing X*^0^ *from Y* It is worth emphasizing that the aforementioned estimation strategy does not restrict compositional factors to only reflect biology-induced global abundance changes; in reality, if feature-wise perturbations (*v*_*gi*_) are also of technical origin, they can well be correlated with other sources of technical variation, and can be seen to estimate technical variation beyond what is accounted for by sample depth adjustments. Thus, it is interesting to ask under what conditions compositional factors arising from scaling techniques (including our proposed technique in this work) can reconstruct *X*^0^. In the supplementary, we show that in the presence of sequence-able experimentally introduced contaminants, utilizing existing compositional correction tools amounts to applying stricter assumptions than the often-cited assumption of “technical biases affecting all feature the same way”. The precise condition is given in the supplement (supplementary section 3, eqn. 6). In the absence of contamination, we find the traditional assumption to be sufficient.

### Existing techniques fail to correct for compositional bias in sparse 16S survey data

In this subsection, we ask how existing techniques fare in estimating compositional correction factors, both in settings at large sample depths and with particular relevance to sparse 16S count data. We will find that library size/subsampling approaches are provably bad and that other scaling techniques face certain difficulties with sparse data. We will also note that the common strategy of deriving normalization factors/data transformations after adding pseudocounts to the original sparse count data transformations also lead to biased estimates of scale factors.

Our analysis below is limited to methods that provide interpretable estimates of fold-changes. We therefore do not consider differential abundance inferences arising from rank-based methods. We also leave the analysis of non-linear normalization techniques for future work.

*Library size/Subsampling based approaches* To understand the practical importance of resolving confounding caused by compositional bias, we first asked under what conditions, inferences made without compositional correction would continue to reflect changes in absolute abundances in an unbiased manner. We formally analyzed its influence within the framework of generalized linear models, a widely used statistical framework within several count data packages (supplementary section 1). Under the most natural adjustments based on the total count (e.g., unaltered R/FPKM/CPM/subsampling/rarefication based approaches), we found that these conditions can be precisely characterized and are extremely limited in their applicability in general experimental settings. It may be tempting to argue that one can resort to total count-based normalization if total feature content is the same across conditions. However, as shown in supplementary section 1, it is easy to see that this assumption is only valid when strict constraints on the levels of technical perturbation of feature abundances and sequence-able contaminants are respected, an assumption that can be very easily violated in metagenomic experiments [36, 37, 38], which usually feature high intra-and inter-group feature diversity.

*Reference normalization and robust fold-change estimation techniques* We now compare and contrast library size adjustments with a few reference based techniques (reviewed in **Table. 1**) in overcoming compositional bias at *high* sample depths. Furthermore, many widely used genomic differential abundance testing toolkits enforce prior assumptions on reconstructed fold changes, and moderate their estimation. This made us wonder about the robustness of these testing techniques in overcoming the false positives that would otherwise be created without compositional bias correction. With an exhaustive set of simulations at high coverage sample depths (similar to bulk RNAseq) with 20*M* reads per sample, by and large, we found that all testing packages behaved the same way, and the key ingredient to overcome compositional bias always was an appropriate normalization technique (supplementary section 2). We also found that reference based normalization procedures out-performed library size based techniques significantly, re-emphasizing the analytic insights we mentioned previously. With sparse 16S data however, such techniques developed for bulk RNAseq faced major difficulties as illustrated next.

In **Fig. 4**, we plot the feature-wise compositional scale estimates (i.e., ratio of sample proportion to that of the reference; third column entries in **Table. 1**), obtained from TMM and DESeq for a sample in two different 16S microbiome datasets. TMM computes a weighted average over these feature-wise estimates, while DESeq proposes the median. The first column corresponds to a bulk RNAseq study of the rat body map [39]; the second corresponds to those from a 16S metagenomic dataset [40]. Strikingly, while a large number of features agree on their scale factors for a sample arising from bulk RNAseq for both TMM and DESeq strategies, the sparse nature of metagenomic count data makes robust estimation of their scale factors extremely difficult. Furthermore, large variance is also observed across the scale factors suggested by the individual features. Clearly, a moderated estimation procedure is warranted.

**Figure 4.**
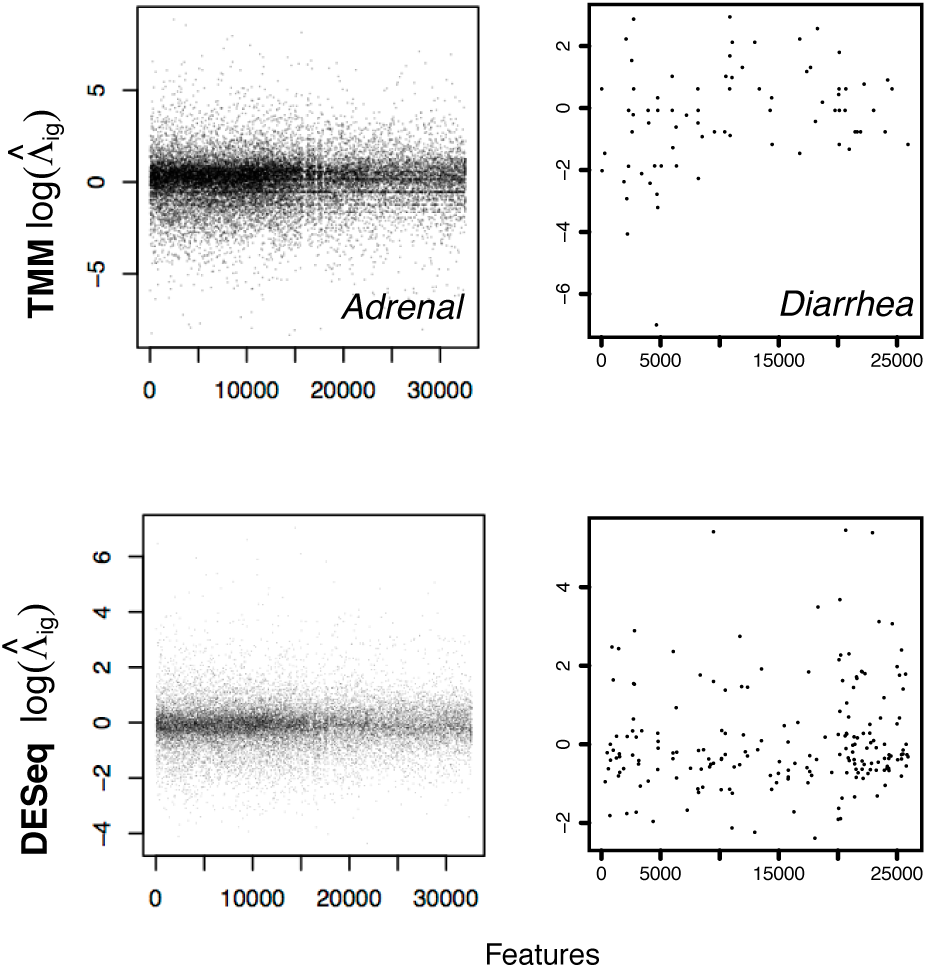
Estimation of compositional correction scales from sparse count data. On the left column, we plot the feature-wise ratio (Λ_*gji*_) estimates adjusted for sample depth from each feature *i* in one of the samples from the Adrenal tissue of the rat body map dataset (bulk RNAseq), and on the right column, we plot the same values arising from a sample in the Diarrheal dataset (16S metagenomics). The top and bottom rows correspond to the scales estimated using TMM and DESeq respectively. In the case of bulk RNAseq data, large numbers of individual feature estimates agree on a compositional scale factor. Simple averaging, or some robust averaging would help us obtain the scale factor exactly. A similar robust behavior is observed with all the tissues available in the bodymapRat dataset (considered later in text). On the second column, we plot the feature-wise ratio values from a metagenomic 16S marker gene survey of infant gut microbiota. There is no general agreement among the features on the scale factors, and simple averaging will not work. We note that what we have shown are fairly good cases. Several samples entertain only a few tens of shared species with an arbitrary reference sample within the dataset. In this work, we aimed to model this variability and estimate the scale factors robustly by borrowing information across features and samples.

One might wonder if adding pseudocounts to the orginal count data (a common procedure in metagenomic data analysis [19, 41]) effectively deals away with the problem. However, as shown in **Fig. 5**, with large number of features absent per sample, these scale factors roughly reflect the value of the pseudocount, and are systematically scaled down in value as sequencing depth, which is strongly correlated with feature presence, increases. This result suggests that addition of pseudocounts to data need not be the right strategy for deriving normalization scales based on CLR [42] or other similar methods, especially when the data is sparse. The alternate idea of only deriving scale factors based on positive values alone, are also associated with problems as we will see later in the text.

**Figure 5.**
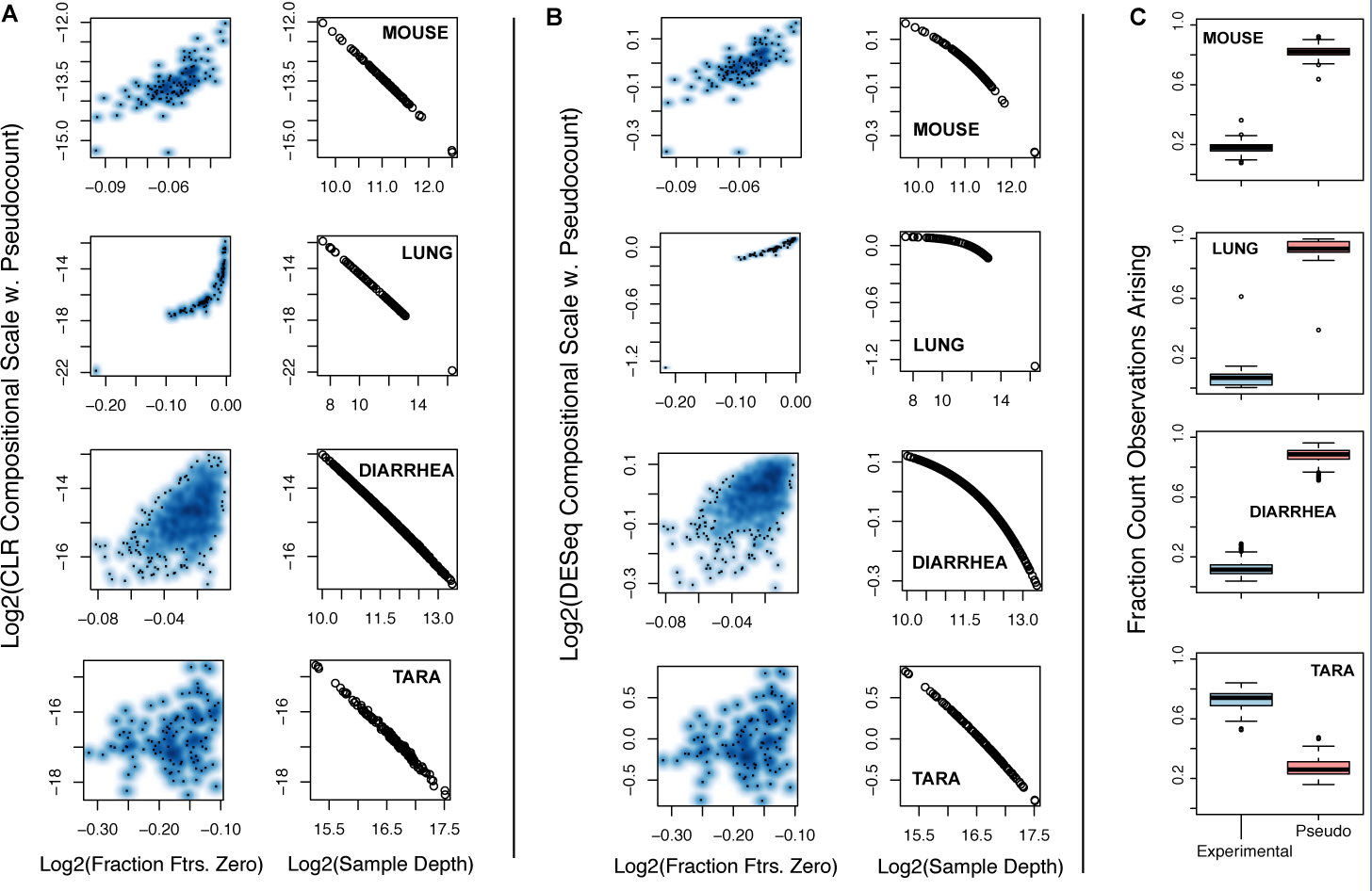
Adding pseudocounts leads to biased normalization. For each of the four microbiome count datasets (rows: Mouse, Lung, Diarrheal and Tara Oceans), we plot (**A**) CLR and (**B**) DESeq compositional scales obtained after adding a pseudo count value of 1, as a function of fraction of features that are zero in the samples (first column) and the sample depth (second column). The observed behavior was not sensitive to the value of pseudocount used. Refer supplementary Fig. 7 for the same plot for a pseudocount value of 10^−7^. (**C**) shows the total number of pseudocounts added, which is essentially the number of features observed in a dataset, and the total actual counts observed in the dataset divided by their sum i.e., the total implied sequencing depth after pseudocounts addition. A large fraction of sequencing depth in the new pseudocounted dataset is now arising from pseudocounts than the true experimental counts, when the data is excessively sparse. Indeed, if the pseudocount value is altered to a very low positive fraction value, the boxplots will reflect reversed locations, but this plot is only used to stress the level of alteration made to a dataset. Only in the Tara Oceans project, where the sample depth is 100K reads, do the boxplots shift. However, at a roughly median 90% features absent, that data when altered by pseudocounts, also leads to biased scaling factors as seen in (A) and (B).

### Our proposed approach (Wrench) reconstructs precise group-wise estimates, and achieves significantly better simulation performance

To overcome the issues faced by existing techniques, we devised an approach based on the following observations and assumptions. First, group/condition-wise feature count distributions are less noisy than sample-wise feature count distributions, and it may be useful to Bayes-shrink sample-wise estimators towards that of group-wise global estimates. Second, zero abundance values in metagenomic samples are predominantly caused by competition effects induced by sequencing technology (illustrated in **Fig. 1**), and therefore can be indicative of large changes in underlying compositions^[1]^ with respect to a chosen reference. Indeed, ignoring sterile/control samples, the median fraction of features recording a zero count across samples in the mouse, lung, diarrheal, human microbiome project and (the very high coverage) Tara oceans datasets were: .96, .98, .98, .98 and .88. These respectively had median sample depths of roughly 2.2*K*, 4.5*K*, 3.3*K*, 4.4*K* and 100*K* reads. In direct contrast, this value for the high coverage bulkRNAseq rat body map across 11 organs at a median sample depth of 9.7*M* reads, is .33. Large number of features, extreme diversity, and time-dependent dynamic fluctuations in microbial abundances can result in such high sparsity levels in metagenomic datasets. When working within the fundamental assumption that *most features do not change across conditions*, such extraordinary sparsity levels can then be attributed, by and large, to competition among features for being sequenced. As we illustrate in **Fig. 6**, zero observations in a sample are correlated with compositional changes, and truncated analyses that ignore them (as is done with TMM / DESeq / metagenomic CSS normalization techniques) effectively leads to loss of information and results that are opposite to what is expected.

**Figure 6.**
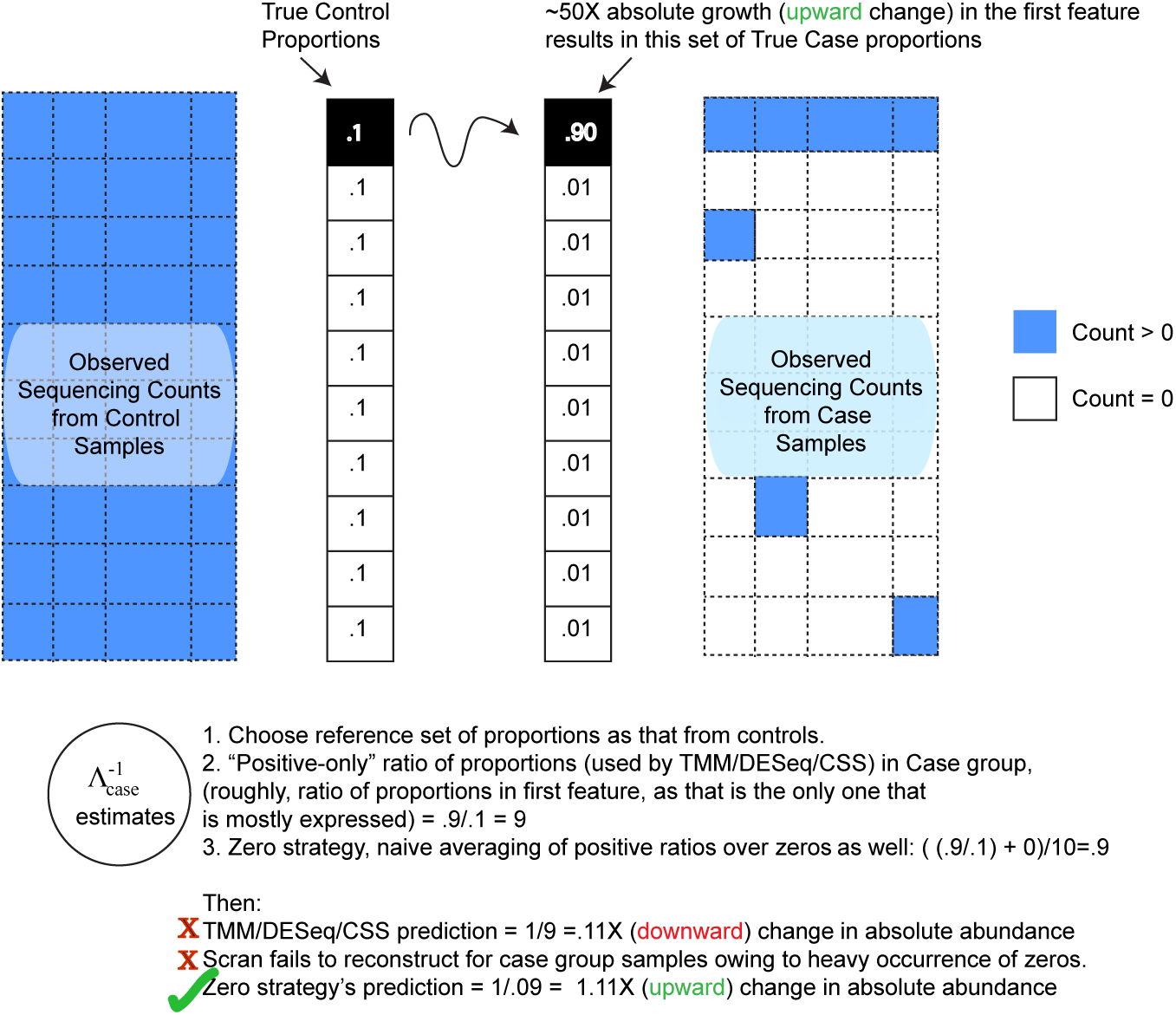
Ignoring zeroes can introduce bias in normalization, when zeroes predominantly arise from under-sampling. An artificial example with 10 features and two groups (“controls” and “cases”), when one of the features undergoes a roughly 50X expansion (a log_2_ fold change of 5.64) in cases compared to controls. This drives the relative abundances of the rest of the 9 features relatively low in the case group. As a result features that are largely present in the controls are not observed in the case group at moderate sequencing depths. Scaling normalization strategies that derive scales based only on the positive count values, can underestimate compositional changes as shown.

We now give a brief overview of the technique (Wrench) proposed in this work. More details are presented in the Methods section. With average proportions across a dataset as our reference, we model our feature-wise proportion ratios as a hurdle log-normal model^[2]^, with feature-specific zero-generation probabilities, means and variances. The analytical tractability of the model allows us to standardize the feature-wise values within and across samples, and derive the compositional scale estimates by basing heavy weights on less variable features that are more likely to occur across samples in a dataset. In addition, to make the computed factors robust to low sequencing depths and low abundant features, we employ an empirical Bayes strategy that smooths the feature-wise estimates across samples before deriving the sample-wise factors. Such situations are rather common in metagenomics, and some robustness to overcome heavy sampling variations is desirable.

**Table. 2** succinctly illustrates where current state of the art fails, while more comprehensive simulations illustrating the effectiveness of the proposed approach is presented in **Fig. 7**. To generate table 2, roughly, we simulated two experimental groups, with 54*K* features whose proportions were chosen from the lung microbiome data, and let 35% of features change across conditions (see Methods for details on simulations). The net true compositional change resulting from each simulation, and their corresponding reconstructions by the various techniques when the count data are generated at different sequencing depths are shown. The following observations form the theme of these, and the more elaborate simulations summarized in Fig. 7: 1)TMM/CSS, because they focus on positive-valued observations only, are restricted in the range of scales they can reconstruct. 2) Scran can yield accurate estimators at very large sequencing depths when high feature-wise coverages are achieved. Unfortunately, this behavior is highly dependent on the underlying feature proportions and their diversity. 3) Wrench estimators offer better alternatives for under-sampled data, and as we shall observe below in their empirical performances, they can still offer robust protection against compositional bias at higher coverages. For specific comparisons with pseudocounted CLR, please refer supplementary Fig. 9, in which we show the proposed technique (Wrench) performing significantly better.

**Table 2.** Example simulations illustrate the limitations of current techniques. Shown are the group-wise true and reconstructed compositional scales from the methods compared on two simulated examples, each at different sequencing depths and at different total true absolute abundance changes for a roughly 54*K* features with control group proportions derived from the Lung microbiome. Low-coverage and/or high compositional changes are problematic for current techniques due to the sparsity they cause in the count data. *W*_1_,… *W*_3_ are Wrench estimators proposed in the Methods section that adjust the base estimator *W*_0_ for feature-wise zero-generation properties. All are presented here for comparison purposes. Our default estimator is *W*_2_.

| Net Compositional Change ($\Lambda_g$) | Average Sample Depth | CLR | TMM | CSS | Scran | $W_0$ | $W_1$ | $W_2$ | $W_3$ |
| --- | --- | --- | --- | --- | --- | --- | --- | --- | --- |
| 36.86X | 1M | 1.36 | 1.45 | 5.41 | 22.57 | 19.32 | 31.44 | 30.65 | 32.01 |
| 7.75X | 10K | .95 | 3.05 | 1.47 | 12.08<br>(14/40 samples failed) | 5.30 | 6.32 | 6.31 | 6.70 |

**Figure 7.**
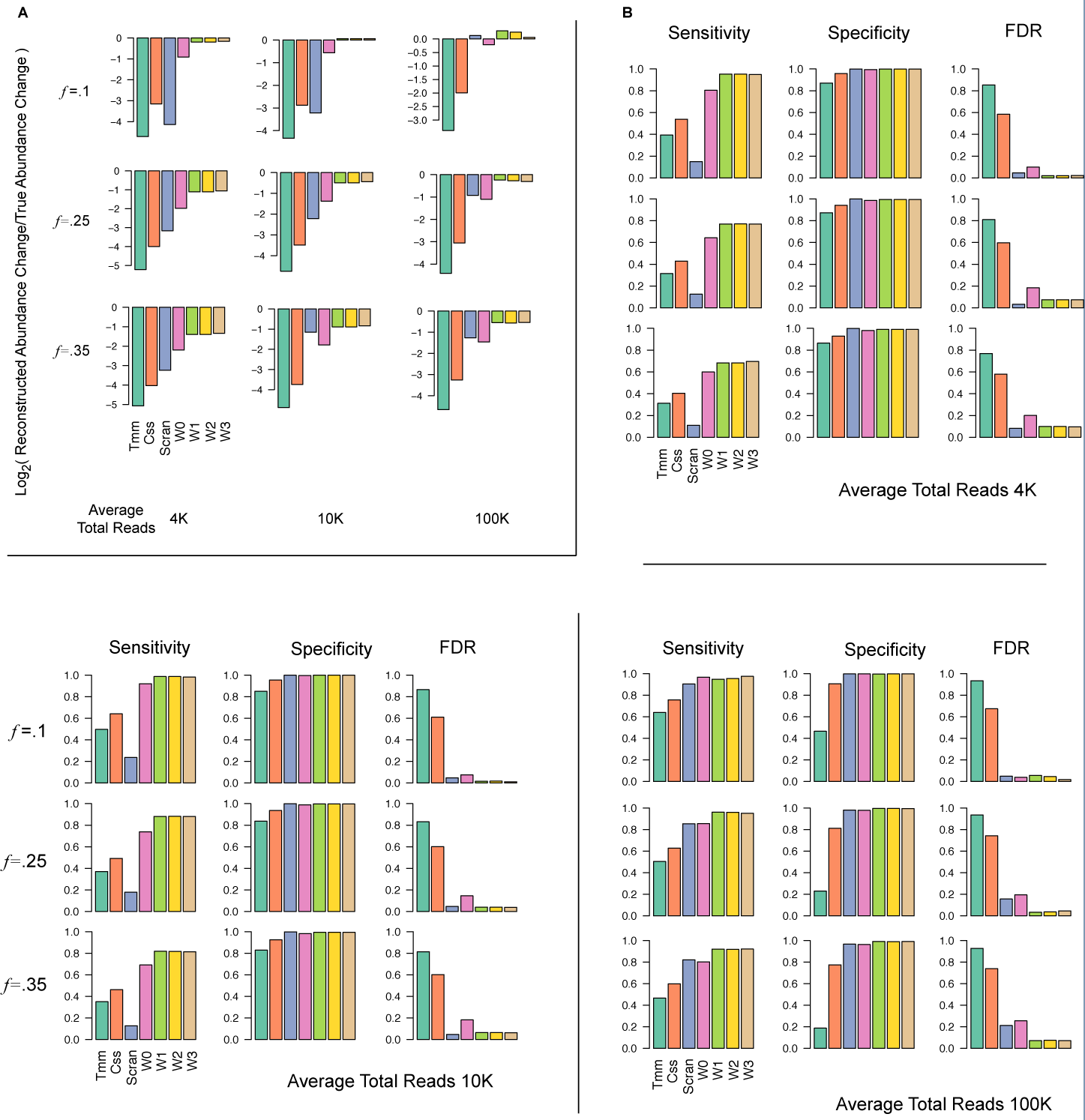
Wrench scales outperform competing approaches in reconstructing compositional changes and in differential abundance testing. Multiple iterations of two group simulations are simulated with various fractions of features perturbed across conditions (rows, *ƒ* in figures), total number of reads. Their average accuracy metrics in reconstruction and differential abundance testing are plotted. The control proportions were set to those obtained from the mouse microbiome dataset. (A) Average log ratios of reconstructed to true absolute abundance changes. Each row corresponds to a particular setting of *ƒ*, and each column a particular setting of average sequencing depth. Scran also suffered from being unable to provide scales for samples in each simulation set (sometimes as high as 60% of the samples at 4K and 10K average reads). (B) Average sensitivity, specificity and false discoveries at FDR .1 of detecting true differential absolute abundances. *W*_0_ is the regularized Wrench estimator without sparsity adjustments and *W*_1_, ‥*W*_3_ are various adjusted estimators compared here. For details on this and simulations, see Methods. Behavior was similar for other parameteric variations (variances of global and sample-wise fold change distributions, number of samples) of simulations.

We briefly note a key ingredient about our simulation procedure. Simulating sequencing count data as *independent* Poissons / Negative Binomials – as is commonly done in benchmarking pipelines – does not inject compositional bias into simulated data. From the perspective of performance comparisons for compositional correction, doing so is therefore inappropriate. A renormalization procedure after assigning feature-wise fold-changes is necessary. Alternatively, if absolute abundances are generated, subsampling to a desired sample depth needs to be performed.

### Wrench has better normalization accuracy in experimental data

Below, we show five different results illustrating the improvements Wrench offers over existing techniques in experimental data. The first two show that Wrench leads to reduced false positive calls in differential abundance inference, while the other three demonstrate the improved quality of positive associations.

*Reduction of false positives* We used two approaches to compare the performance of Wrench in reducing false positive calls in differential abundance inference. Each of these analyses was performed across all biological groups with atleast 15 samples in the mouse (2 diet types), Diarrheal (2 groups), Tara (5 oceans), HMP (JCVI, 16 body sites), and HMP (BCM, 16 body sites) and averaged the results across these 41 experimental groups.

We ignored the lung microbiome for these analyses as Scran had particular difficulty making direct comparisons hard. Owing to the heavy sparsity in these datasets, Scran failed to provide scales for 53 out of 72 samples of the lung microbiome, 10 out of 132 observations of the mouse microbiome, 6 out of 992 samples of the diarrheal dataset. Notice that Wrench not only recovers compositional scales for these samples, but also at magnitudes that were coherent with other samples from similar experimental groups (see next subsection) indicating some validity for the computed normalization factors.

First, a standard resampling analysis was performed. For every given experimental group, two artificial groups are repeatedly constructed via resampling (without replacement), and the total number of significant calls made during differential abundance analysis is recorded in each repetition. For each iterate, we compute the log_2_(*F*_*Other*_/*F*_*Wrench*_) ratio, where *F*_*Other*_ is the total number of significant calls made by a competing method (Total Sum / TMM / Scran / CSS) and *F*_*Wrench*_ is the total number of significant calls made by Wrench. If Wrench is superior these logged ratios should be > 0. The average of these ratios across all the experimental groups mentioned above is plotted in **Fig. 8A**, and we find Wrench meeting the goal. Although total sum does not show a significant difference in this analysis, as illustrated next, it is insufficient in capturing the null variation in the data.

**Figure 8.**
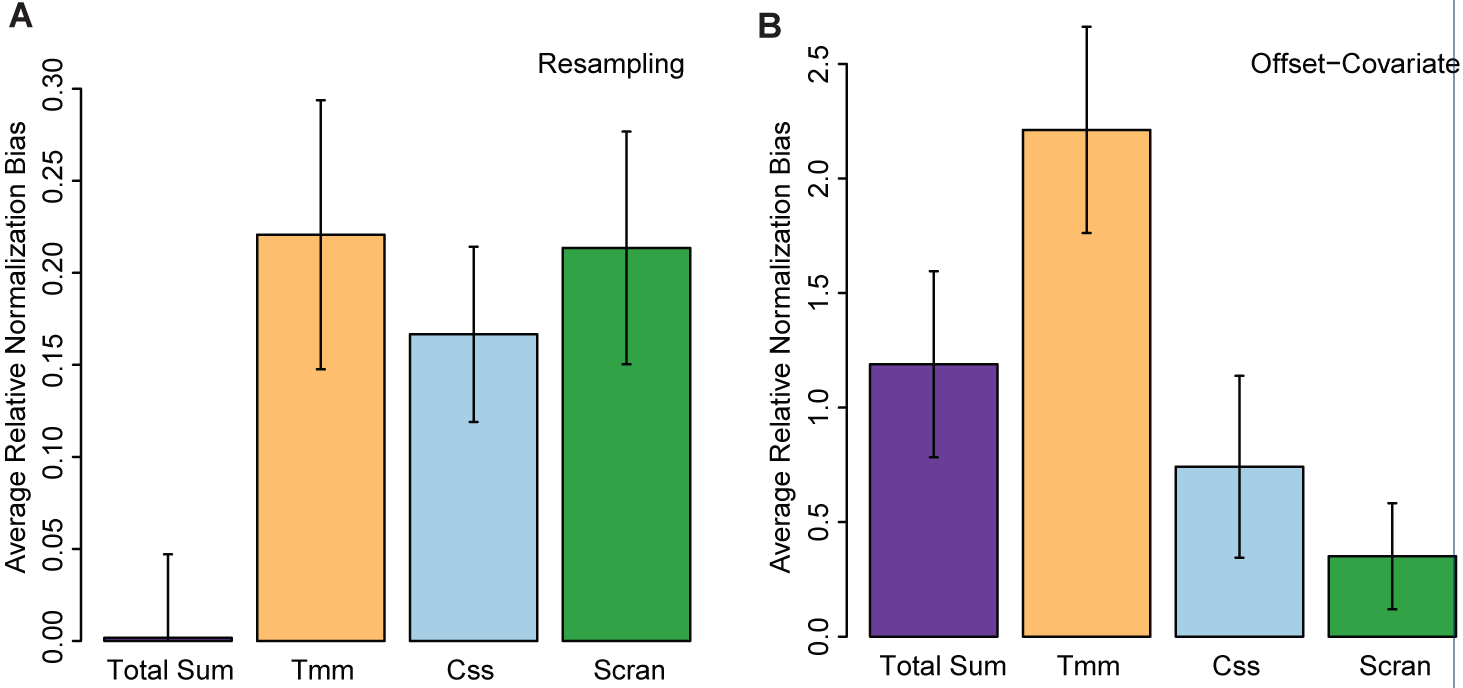
Wrench scales lead to reduced false positive calls. (A) The average of log_2_(*F*_*Other*_/*F*_*Wrench*_) values obtained over artificial two group splits of homogeneous experimental group data is shown and (B) the average of log_2_(*C*_*Other*_/*C*_*Wrench*_) values across 41 metagenomic experimental groups are shown. Standard error bars are shown. In both plots, positive values for a method imply reduced accuracy relative to Wrench. *F*_*Other*_: total number of diffferentially abundant features found by a competing method (total sum, TMM, CSS or Scran). *F*_*Wrench*_: total number of differentially abundant features found by Wrench. *C*_*Other*_: total number of features where the covariate term for Wrench normalization factors were found to be significant when competing method is used as offset. *C*_*Wrench*_: total number of features where the covariate term for a competing method’s normalization factors were found to be significant, when Wrench is used as covariate.

We next exploited the offset-covariate approach introduced in [25]. For every feature/OTU within a homogenous experimental group, two generalized linear models are fitted: in model (a) Wrench normalization factors as offset, and those of a competing method as covariate. In model (b), normalization factors from a competing method as offset, and those of Wrench as covariate. The number of features for which the covariate term was called significant is recorded in both (a) and (b). We will denote them respectively as *C*_*Wrench*_ and *C*_*Other*_. If Wrench sufficiently captures the variation in data, the number of times the covariate term from a competing method is called significant will be low. That is: the logged ratio log_2_(*C*_*Other*_/*C*_*Wrench*_) must be > 0. The average of these values across all the experimental groups mentioned above is plotted in **Fig. 8B**, and we find Wrench to improve upon other techniques.

*Improved association discoveries* To compare the quality of associations achieved with the various normalization methods, we re-analyzed the Tara Oceans 16S microbiome dataset.

Even though the contribution of true compositional changes and other technical biases are not identifiable from the compositional scales without extra information, we asked if the reconstructed scales correlate with orthogonal information on absolute abundances, and other measures of technical biases. The results are summarized in **Table 3**. Interestingly, in the very high coverage Tara Oceans metagenomics project, Wrench and Scran estimators achieve comparable correlations (>50%) with absolute flow cytometry measurements of microbial counts from the Tara Oceans project. Scran failed to reconstruct the scales for 3 samples. TMM and CSS had substantially poor correlations. Similarly, Wrench normalization factors had comparable/slightly better correlations to the total ERCC spike-in counts in bulk and single cell RNAseq datasets. In direct contrast, CLR scale factors (the geometric means of proportions) computed with pseudocounts were either uncorrelated or highly anti-correlated with the aforementioned measurements reflecting technical biases. These results reaffirm that there are advantages to exploiting specialized compositional correction tools even with microbiome datasets teeming with microbes of extraordinary diversity.

**Table 3.** Correlations of compositional scales with orthogonal measurements on absolute abundances/technical biases. Correlations of logged reconstructed abundance factors (1/compositional correction factor) with logged total flow cytometry cell counts is shown for the Tara project. Correlations of logged normalization factors with logged total ERCC counts are shown in the case of the rat body map and embryonic stem cells datasets. Given the high sparsity in these datsets, CLR factors computed by adding pseudocounts, essentially had no information on technical biases. *W*_1_, … *W*_3_ are estimators proposed in the Methods section that adjust the base estimator *W*_0_ for feature-wise zero-generation properties. All are presented here for comparison purposes. The default Wrench estimator (*W*_2_) compares well at low and high coverage settings. For more details on these and the distinction in terminology between compositional correction factors and normalization factors, refer Materials and Methods.

| Dataset | Type | CLR | TMM | CSS | Scran | $W_0$ | $W_1$ | $W_2$ | $W_3$ |
| --- | --- | --- | --- | --- | --- | --- | --- | --- | --- |
| Tara Oceans [10] | 16s (from Whole Metagenome) | 0 ($-2.65 \times 10^{-6}$) | 0.26 | 0.15 | 0.52 | .58 | .54 | .53 | .53 |
| Rat BodyMap [39] | Bulk RNAseq | -0.36 | 0.22 | 0.16 | 0.18 | .20 | .19 | .20 | .26 |
| Embryonic Stem Cells [53] | UMI/scRNAseq | -0.70 | .70 | .67 | .67 | .71 | .70 | .70 | .68 |

We next analyzed the quality of differential abundance inference arising from competing normalization techniques, by performing two sets of enrichment analyses.

In the first procedure, we extracted broad genus-level functional annotations from the Faprotax database [43], and tested for their enrichment in positively associated genera in the deep chlorophyll (DCM) and the mesopelagic layer (MES) samples of the oceans relative to the surface layer. The total number of significantly differentially abundant OTU calls were widely different across techniques: Wrench and Scran made roughly 30% fewer calls compared to total sum, TMM, and CSS. Given the relatively general nature of the annotations, all methods yielded expected annotations in the DCM and MES layers based on previous studies, although there were a few differences (additional file 2). Nitrite respiration/reduction/anoxygenic pho-totropy, oil bioremediation were found enriched in mesopelagic layer by all methods, while methanogenesis, a function that is usually associated with mesopelagic and deep sea microbes [44, 45, 46, 10, 43] was not found enriched in MES by total sum. Both Wrench and Scran did not find xylanolysis to be enriched in the mesopelagic layer, while other methods did. We were unable to find literature evidence supporting this call, and the result could potentially be due to the higher number of OTUs called differentially abundant by the other methods. Aerobic ammonia/nitrite oxidation and fixation were found to be enriched in DCM by all methods. Total sum and TMM found a methanogenesis related module enriched in DCM, while other methods did not.

To evaluate the methods in a more fine-grained setting, we devised the following validation approach. The design of the Tara oceans experiments - where 16S reconstructions are obtained from whole metagenome shotgun sequencing data - makes the following analysis feasible. Because the Tara project’s functional (gene content summarized as Kegg Modules, KMs) and 16S data arise from the same input DNA samples, the same compositional factors should apply for both datatypes. We therefore estimated compositional factors from 16S data using the different normalization methods and applied the resulting estimates to the KM abundance data from the corresponding matched samples. Next, we computed Spearman rank correlation between OTU and KM normalized abundances and annotated OTUs with those KMs which showed correlation of at least 0.75. Finally, we identified OTUs that were positively associated with each layer using differential abundance analysis. With the KM annotations in place, we performed Fisher exact tests to compute the enrichment scores in the identified OTUs. Detailed tables are provided in *additional file 2*. In mesopelagic samples, Scran finds enrichment in only 30 KMs, while other methods recovered at least 100 KMs. Specifically, ureolysis, motility, several denitrification/methanogenesis processes and aminoacid biosynthetic/transport mechanisms (functions that have been attributed to microbes in the mesopelagic layer and deep sea) [47, 48, 10, 43], were missed by Scran, while Wrench finds them. On the other hand, Total sum, TMM and CSS found more varied and general processes including various ribosomal, transcription/translation components to be enriched in both MES and DCM layers.

Notice that the first analysis gives a broad sense of the genera identified by the competing methods in light of existing annotations, while the second gives a sense of the quality of annotations one might confer on the OTUs based on the normalized expression levels of OTUs and the measured functional content themselves. In both cases, Wrench is shown to retain relevant information, and the relatively more specific nature of the latter analysis reveals that Wrench demonstrably improves upon other methods.

#### Inferences following compositional correction show improved coherence with experimental data

We further demonstrate the impact of compositional bias in downstream inference below. The experimental cell density measurements in the Tara Oceans project show a highly significant overall reduction in the mesopelagic samples when compared the surface layer (see Fig. 3 in ref [10]). Thus, we expect an overall negative change in the reconstructed fold changes, when performing a differential abundance analysis of the OTUs across these two ocean layers.

Summing the log-fold changes of significantly associated OTUs (both positive and negative) serves as a measure of a net change experienced by a community. If a given method produces fold change inferences that track the above mentioned empirical cell density measurements, we expect it to yield an overall negative net change value for the significantly differentially abundant OTUs in the mesopelagic community. As illustrated in **Fig. 9A**, this value for total sum normalized data is +10577.99, while that for Wrench is –8919.65, showing that differential abundances arising from Wrench agrees more appropriately with the underlying community change. **Fig. 9B and C**, show how these values distribute across the major phyla focussed in the Tara oceans article. These plots demonstrate that the two approaches lead to markedly different conclusions on the net change experienced by a phylum. In particular, Proteobacteria, Actinobacteria, Euryarchaeota were predicted to have drastically high positive changes by total sum (while Wrench predicts a marked decrease in the negative direction), and sizable differences were apparent in the values obtained with the rest of the phyla.

**Figure 9.**
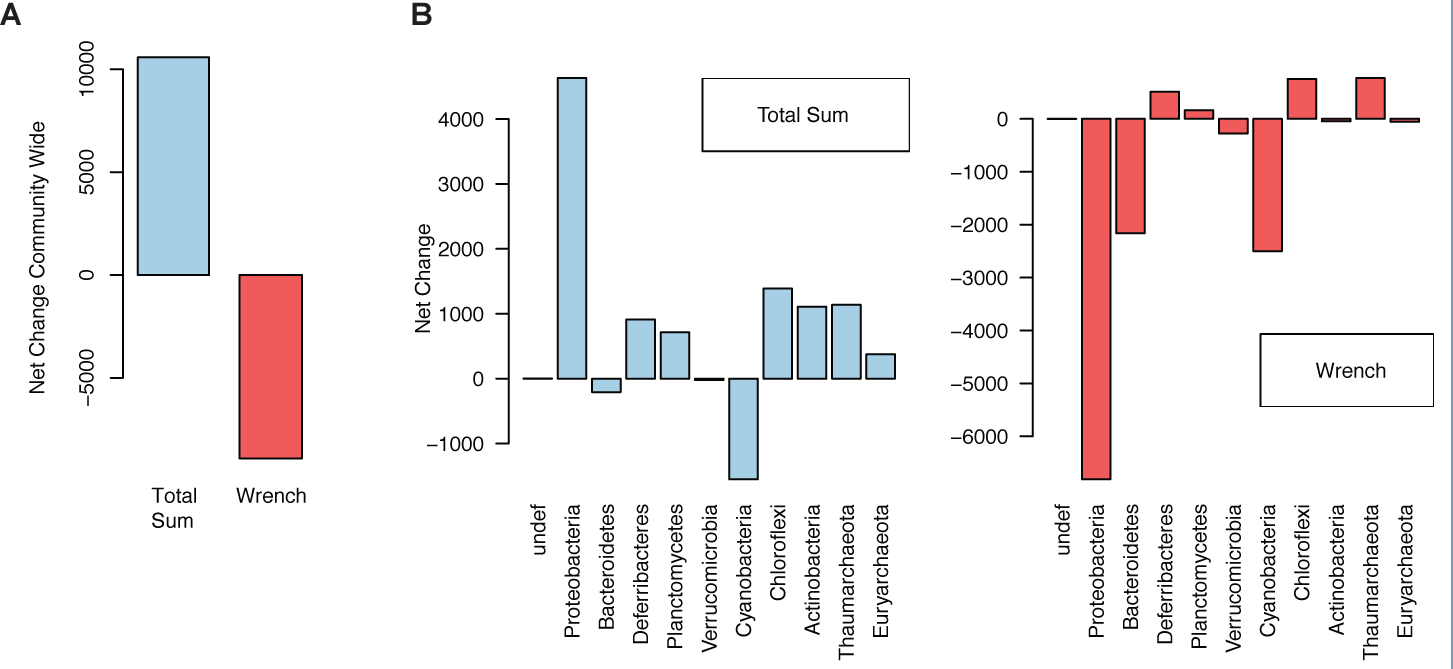
Wrench normalized data lead to better downstream inferences. (A) The sum of log-fold changes of differentially abundant OTUs is used as a measure of net change experienced by a community. This value is plotted for the differentially abundant OTUs in the mesopelagic ocean layer relative to the surface layer in the Tara oceans 16S data, for Total Sum and Wrench normalization. (B) The same metric plotted for various major phyla of interest in the Tara oceans project.

### Compositional scale factor estimates imply substantial technical biases, indicating importance of further experimental studies

We next analyzed the phenotypic integrity of the compositional scales reconstructed by the various methods. In the absence of technical biases, following our discussion in the previous subsection, compositional factors should hover around 1 (upto some arbitrary scaling). This is *not* what we observe in samples from metagenomic datasets. All scale normalization techniques resulted in group-wise integrity in the scales they reconstructed within and across related phenotypic categories, potentially indicating the general importance of correcting for confounding induced by compositional bias in general practice. Total sum normalization is oblivious to these biases, making further experimental studies on compositional bias important. For instance, in the microbiome samples arising from the Human Microbiome Project, as shown in **Fig. 10A**, we noted systematic body site-specific global deviations in the fold change distributions. This is similar to what was illustrated with the Tara project in Fig. 2. We found the reconstructed compositional scales to largely organize by body sites, across normalization techniques (**Fig. 10B**), behind-ear and stool samples were distinctly located in terms of their compositional scales from the oral and vaginal microbiomes (notice the log scale in these plots). This behavior was also recapitulated in scales reconstructed from other centers. Supplementary Figs. 10 and 11 present similar results on samples arising from the J. Craig Venter Institute. In the case of the mouse microbiome samples, most normalization techniques predicted a mild change in differential feature content across the two diet groups (**Fig. 10C**, and supplementary Fig. 12). In the lung microbiome, the lung and oral cavities had roughly similar scales across smokers and non-smokers (supplementary Fig. 13), while scales from the probing instruments had relatively higher variability, which we found to directly correlate with the high variability of feature presence in the count data arising from these samples. In the diarrheal datasets of children, however, no significant compositional differences were found across the various country/health-status populations (**Fig. 10D**).

**Figure 10.**
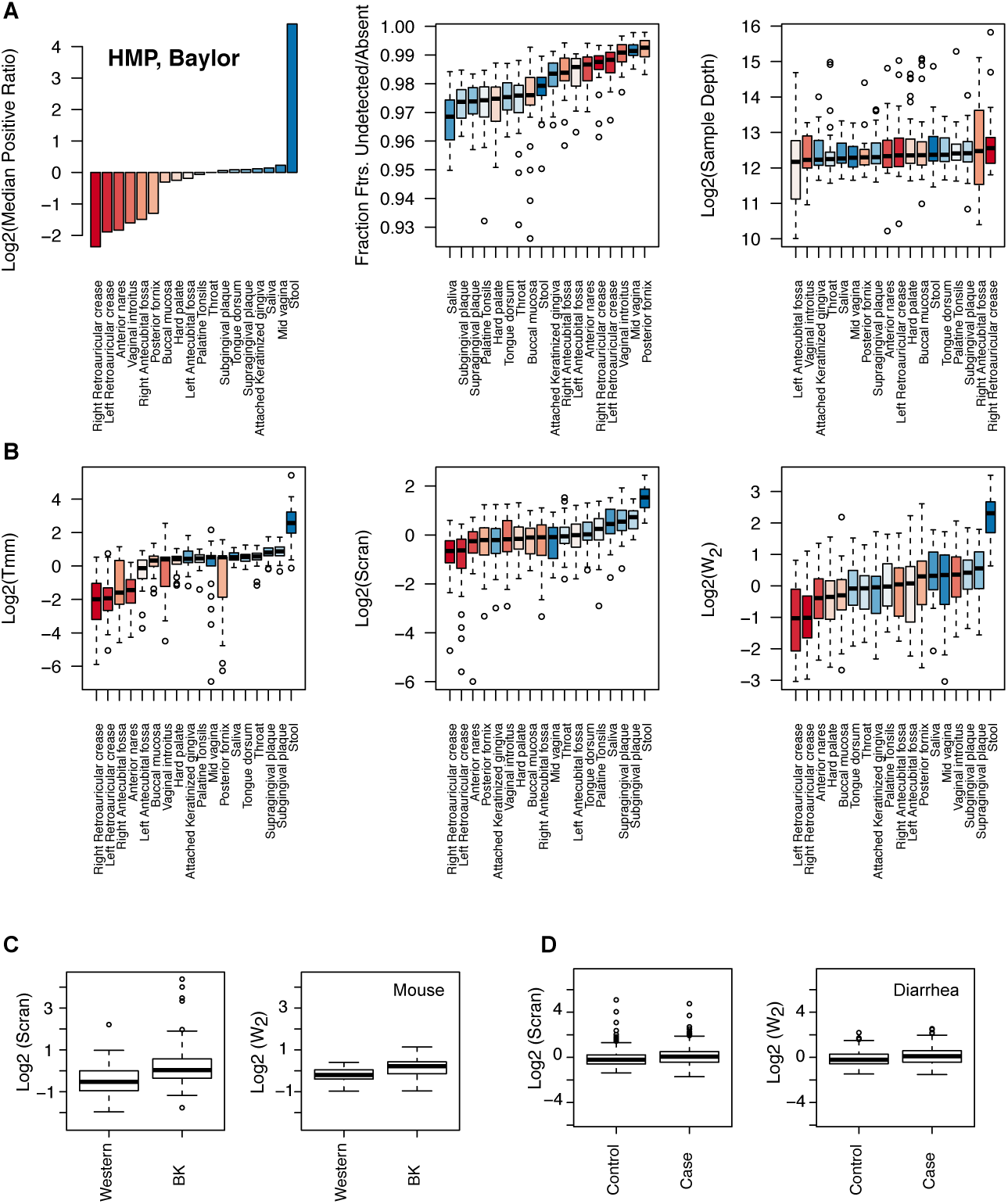
Wrench retains potential biological information, and indicates importance of compositional correction in general practice. We plot some statistical summaries and the compositional scale factors reconstructed by a few techniques for various Human Microbiome Project samples, sequenced at the Baylor College of Medicine. (**A**) On the top-left, we plot the logged median of the positive ratios of group-averaged proportions to that of *Throat* chosen as the reference group. Stool samples show considerable deviation from the rest of the samples despite having comparable fraction of features detected and sample depths to other body sites. Notice the log scale. (**B**) The similarity in the reconstructed scales across techniques (second row) for closely related body sites are striking; although minor variations in the relative placements were observed across centers potentially due to technical sources of variation, the overall behavior of highly significant differences in the scales of behind-ear and stool samples were similar across sequencing centers ( supplementary Fig. 10) and normalization methods. Corresponding CSS scales in supplementary Fig. 11. These techniques predict a roughly 4X-8X (ratio of medians)inflation in the Log2-fold changes when comparing abundances across these two body sites. (**C**) Wrench and scran compositional scale factors across the plant-based diet (BK) and Western diet (Western) mice gut microbiome samples. (**D**) Compositional scale factors for healthy (Control) and diarrhea afflicted (Case) children. Slight differences in the compositional scales are predicted in the diet comparisons with t-test p-values < 1e-3 for all methods except TMM, but not as much in the diarrheal samples.

For completeness, we also attach similar results from all the 11 organs of the rat body map dataset in the supplementary Fig. 15.

## Discussions

For some researchers, statistical inference of differential abundance is a question of differences in relative abundances; for others, it is a matter of characterizing differences in absolute abundances of features expressed in samples across conditions [49, 14]. In this work, we took the latter view and aimed to characterize the compositional bias injected by sequencing technology on downstream statistical inference of absolute abundances of genomic features.

It is clear that the probability of sequencing a particular feature (ex: mRNA from a given gene or 16S RNA of an unknown microbe) in a sample of interest is not just a function of its own fold change relative to another sample, but inextricably linked to the fold changes of the other features present in the sample in a systematic, statistically non-identifiable manner. Irrevocably, this translates to severely confounding the fold change estimate and the inference thereof resulting from generalized linear models. Because the onus for correcting for compositional bias is transferred to the normalization and testing procedures, we reviewed existing spike-in protocols from the perspective of compositional correction, and analyzed several widely used normalization approaches and differential abundance analysis tools in the context of reasonable simulation settings. In doing so, we also identified problems associated with existing techniques in their applicability to sparse genomic count data like that arising from metagenomics and single cell RNAseq, which lead us to develop a reference based compositional correction tool (Wrench) to achieve the same. Wrench can be broadly viewed as a generalization of TMM [13] for zero-inflated data. We showed that this procedure, by modering feature-wise zero generation, reduces the estimation bias associated with other normalization procedures like TMM/CSS/DESeq that ignore zeroes while computing normalization scales. In addition, by recovering appropriate normalization scales for samples even where current state of the art techniques fail, the method avoids data wastage and potential loss of power during differential expression and other downstream analyses (We catalog a few potential ways by which compositional sources of bias can cause sparsity in metagenomic and single cell sequencing count data in Supplementary section 6). A few important insights emerge.

In our simulations, we found reference based normalization approaches to be far superior in correcting for sequencing technology-induced compositional bias than library size based approaches. From a more practically relevant perspective, we found that in all the tissues from the rat body map bulk RNAseq dataset, the scale factors can be robustly identified. We expect that in other bulk RNAseq datasets, the assumptions underlying compositional correction techniques to hold well. These results reinforce trust in exploiting such scaling practices for other downstream analyses of sequencing count data apart from differential abundance analysis; for example, in estimating pairwise feature correlations. In the regimes where assumptions underlying these techniques are met, an analyst need not be restricted to scientific questions pertaining to relative abundances alone. The fundamental assumption behind all the aforementioned techniques (including our own) is that most features do not change across conditions. As we illustrated, these assumptions appear to hold rather well in bulk RNAseq. Do we expect these to hold in arbitrary microbiome datasets as well? This question is not easy to address without more experiments, but the relatively high correlations obtained with orthogonal measurements of technical biases, the similarity in the compositional scales obtained within samples arising from biological groups, and their sometimes highly significant shifts preserved across normalization techniques and across sequencing centers in large scale studies certainly reinforce the critical importance of characterizing compositional biases, if any, in metagenomic analyses by establishing carefully designed spike-in protocols. In particular, given the inverse dependence of compositional correction factors on the total feature content in the absence of technical biases, the large compositional scale estimates obtained for stool samples (across all normalization techniques) is suspect. Compositional effects can amplify even when a few features experience adverse technical perturbations, and only carefully designed experiments can isolate these effects to inform further normalization approaches. Finally, our results also emphasize the tremendous care one needs to exercise before applying the most natural normalizations based on total sequencing depth or by applying pseudocounts when the data is excessively sparse (CLR, RPKM, CPM, rarefication are a few examples).

This brings us to the question of how effective spike-in strategies are in enabling us to overcome compositional bias. It is immediately clear that the widely used ERCC recommended spike-in procedure for RNAseq cannot help us in overcoming confounded inference due to compositional bias for the simple reason that it already starts with an extract, a compositional data source (supplementary section 2). If one is able to add the spike-in quantities at a prior stage during feature extraction, we would have some hope. Lovén et al., [50] demonstrate a procedure for RNAseq that precisely does this, in which the spike-ins are added at the time when the cells are lysed and suspended in solution [51]. One can perhaps extend these solutions to metagenomics, where we may expect confounding due to compositionality to be heavy by adding barcoded 16S RNAs during feature extraction. We expect similar problems to arise in other genomic and epigenetic measurement techniques that exploit sequencing technology, and the need for the development of appropriate spike-in procedures should be addressed.

Finally, it is imperative that we enforce new tools and techniques for normalization and differential abundance analysis of sequencing count data be benchmarked for compositional bias at least in the simulation pipelines. Data analyses based on large-scale integrations of different data types for predicting clinical phenotypes is increasingly common, and care should be taken to include effective normalization techniques to overcome compositional bias. We hope the results and ideas presented and summarized in our paper enables a researcher to do just that.

## Conclusions

Compositional bias, a linear technical bias, underlying sequencing count data is induced by the sequencing machine. It makes the observed counts reflect relative and not absolute abundances. Normalization based on library size/subsampling techniques cannot resolve this or any other practically relevant technical biases that are uncorrelated with total library size. Reference based techniques developed for normalizing genomic count data thus far, can be viewed to overcome such linear technical biases under reasonable assumptions. However, high resolution surveys like 16S metagenomics are largely undersampled and lead to count data that are filled with zeroes, making existing reference based techniques, with or without pseudocounts, result in biased normalization. This warrants the development of normalization techniques that are robust to heavy sparsity. We have proposed a reference based normalization technique (Wrench) that estimates the overall influence of linear technical biases with significantly improved accuracies by sharing information across samples arising from the same experimental group, and by exploiting statistics based on occurrence and variability of features. Such ideas can also be exploited in projects that integrate data from diverse sources. Results obtained with our and other techniques, suggest that substantial compositional differences can arise in (meta)genomic experiments. Detailed experimental studies that specifically address the influence of compositional bias and other technical sources of variation in metagenomics are needed, and must be encouraged.

## Materials and Methods

### An approach (Wrench) for compositional correction of sparse, genomic count data

Briefly, our normalization strategy can be described as follows. Based on eqn. 1, for a chosen reference vector *q*_0_., accounting for sample depth *τ*_*gj*_, the mean model for the observed positive count of the *i*^*th*^ feature can be written as: 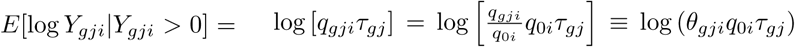, where 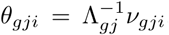. Thus the true ratio of proportions *θ*_*gji*_ encapsulate both the constant 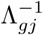 and the absolute fold changes *v*_*gji*_, and can be viewed as the *net* fold change experienced by feature *i* in sample *j* from group *g*. To reflect the assumption that most features do not change across conditions, as is commonly done in genomics, we assume that log *v*_*gji*_ has a zero mean Gaussian distribution. It then follows that log *θ*_*gji*_ follows a Gaussian distribution with a mean parameter 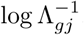. Thus, a robust location estimate of *θ*_*gji*_ for every sample leads us to the desired compositional scale estimate 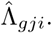. Below, we first illustrate how the *θ*_*gji*_ are estimated, and subsequently discuss the robust averaging procedure.

*Model* We assume the following model for the counts *Y*_*gji*_:

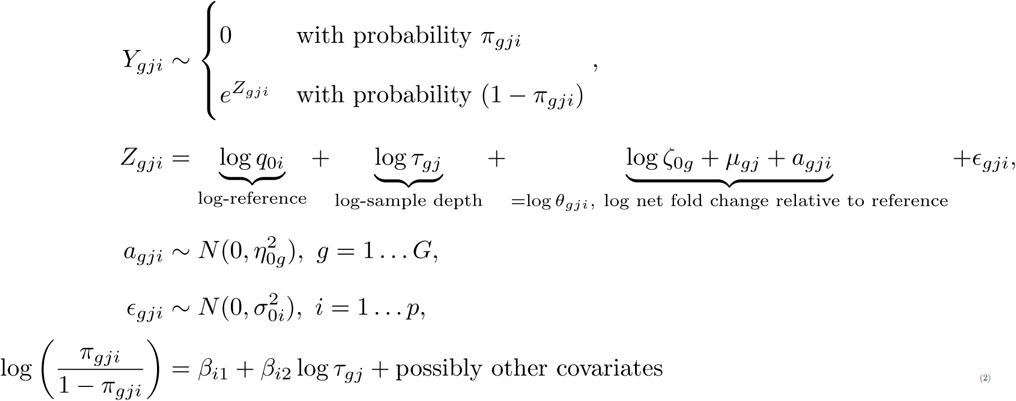

The model assumes the following. For each sample *j* from group *g*, the *i*^*th*^ feature’s count value is sampled from a hurdle log-normal distribution, in which with probability *π*_*gji*_, a value of 0 is realized; and with probability 1 - *π*_*gji*_ a positive count is observed. The probabilities *π*_*gji*_ are determined by sample covariates, including the total sequencing depth. The positive count value is realized as an exponential of a Gaussian random variable *Z*_*gji*_ the mean of which is determined (in accordance with the eqn. 1) by the chosen reference value *q*_0*i*_, sample-depth *τ*_*gj*_, and the *net* fold change 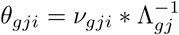, the log of which has been modeled in the above equation as a sum of group-wise effect (log*ζ*_0_*g*__), two-way group-sample interaction (*μ*_*gj*_), a three-way group-sample-feature interaction random effect *a*_*gji*_ and a noise term.

*Estimation of regularized ratios* 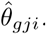: In the model, the 0 subscripted parameters are considered known, and are determined the following way. *τ*_*gj*_ = *Y*_*gj*+_ is the total count of sample *gj*. The reference value for each feature *i*, *q*_0*i*_, is set to the average proportion value 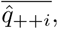, where 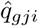 is the observed proportion of feature *i* in sample *gj*, i.e., 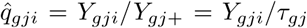. The mean and variance parameters log *ζ*_0_*g*__ and 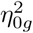 of the Gaussian prior distribution on the log *θ*_*gji*_ are determined based on the corresponding moments of the corresponding empirical distribution of the group-wise pooled raw ratios of proportions: 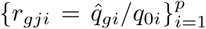. Here, 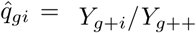 i.e., the overall proportion of feature *i* in the samples from the entire group. Specifically, we fix the group-wise compositional scale 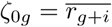 i.e., as the average of the raw ratios including the zero values (following discussions in Fig. 6). We set the variance parameter 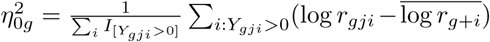 i.e., as the empirical variance of the logged-ratios. Finally, the feature-specific expression variances 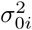 are fixed with values obtained from Limma/Voom. With the above fixed, the unknown parameters *μ*_*gj*_ and *a*_*gji*_ are estimated/predicted using standard random effects estimators: 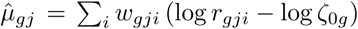 with 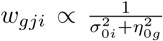, and 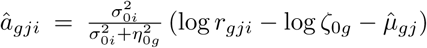. The identifiability of these terms is ensured as the other variance components are fixed. The 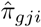 are estimated with logistic regression. The regularized ratios are then calculated as: 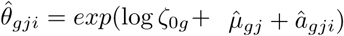.

*Robust averaging of the* 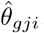: While averaging over the regularized ratios 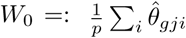 would be one estimation route to 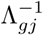, better control can be achieved by taking the variation in the feature-wise zero generation in to account. We shall notice that 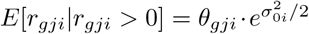, and so a robust averaging over 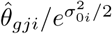, can serve as an estimator of 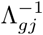. One might choose the weights for averaging to be proportional to that of the inverse hurdle/inclusion probabilities (as is done in survey analysis) 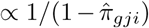 or on the inverse marginal variances ascribed by our model above 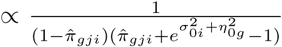. An estimator that we also found to work well empirically is a weighted average of 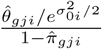 with weights proportional to 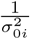. Supplementary section 7 sketches the derivations.

An advantage of these weights (and hence the model) is that the weighting strategies proceed smoothly for features with zero expression values as well, unlike the binomial weights employed in the TMM procedure. Furthermore, when constructing averages, the weights have a favorable property of downweighting zeroes at higher sample depths relative to those in samples at lower sample depths.

In summary, we explored the performance of the following estimators:

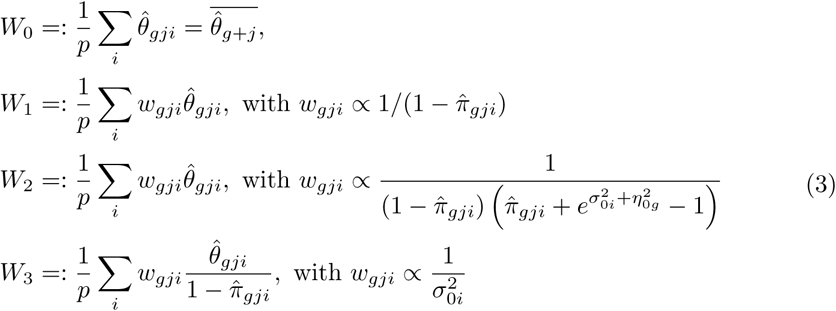

We have found *W*_1_, *W*_2_ and *W*_3_ to work comparably well in simulations and empirical comparisons, and *W*_0_ slightly less so at high sparsity levels at low sample depths. We prefer *W*_2_ as it systematically integrates both the hurdle and positive component variations. In our software implementation, users have the option for other weighted variants, and whether weighted averaging over zeroes is necessary as they see fit. Software documentation and supplementary material embark on further discussions on these ideas.

Finally, with this framework setup, extensions for *batch correction* can be immediately made; this work is being planned for a forthcoming submission.

### Data

We principally demonstrate our results with five datasets from metagenomic surveys. A smoking study (*n* = 72) where the lung microbiome of smokers and non-smokers were surveyed (along with the instruments that were used to sample the individual). A diet study in which the gut microbiomes (*n* = 139) of carefully controlled laboratory mice fed plant-based or western diets were sequenced [32]. A large scale study of human gut microbiomes (*n* = 992) from diarrhea-afflicted and healthy children from various developing countries [40]. 16S metagenomic count data corresponding to all these studies were obtained from the R/Bioconductor package `metagenomeSeq` [26]. The Tara Oceans project’s 16S reconstructions from whole metagenome shotgun sequencing (*n* = 139) was downloaded from The Tara Oceans project website under http://ocean-microbiome.embl.de/data/miTAG.taxonomic.profiles.release.tsv.gz. The flow cy-tometry counts for autotrophs, bacteria, heterotrophs, picoeukaryotes were obtained from TaraSampleInfo_OM.CompanionTables.txt from the same website and summed to serve as a rough measure of total cell count that correlates with sequence-able DNA material. The Human Microbiome Project count data were downloaded from http://downloads.hmpdacc.org/data/HMQCP/otu_table_psn_v35.txt.gz, and the associated metadata are from v35_map_uniquebyPSN.txt.bz2 under the same website.

The processed bulk-RNAseq data corresponding to the rat body map from [39] was obtained from [52].

The UMI single cell RNAseq data from Islam et al., [53] was downladed from GEO under accession GSE46980.

### Implementation of normalization and differential abundance techniques

All analysis and computations were implemented with the R 3.3.0 statistical platform. EdgeR’s `compNormFactors` for TMM, DESeq’s `estimateSizeFactors`, Scran’s `computeSumFactors` (with `positive=TRUE` in sparse datasets) and `metagenomeSeq`’s `calcNormFactors` for CSS were used to compute the respective scales. Implementation of CLR factors used a pseudo-count of 1 following [41], and were computed as the denominator of column 3 in table 1. Limma’s eBayes in combination with lmFit, edgeR’s `estimateDisp, glmFit` and `glmLRT`, DESeq2’s `estimateDispersionsGeneEst` and `nbinomLRT` were used to perform differential abundance testing [54]. Welch’s t-test results were obtained with `t.test`.

### Implementation of Wrench

Wrench is implemented in R, and is available through the `Wrench` package at https://gitlab.umiacs.umd.edu/smuthiah/Wrench.

### Simulations

Given a set of control proportions *q*_1*i*_ for features *i* = 1… *p*, and the fraction of features that are perturbed across the two conditions *ƒ*, we sample the set of true log fold changes (log *v*_*gi*_) from a fold change distribution (fold change distribution) for those randomly chosen features that do change. The fold change distribution is a two-parameter distribution chosen either as a two-parameter Uniform or a Gaussian. Based on the expressions from the first subsection of the results section, the target proportions were then obtained as 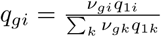. Conditioned on the total number of sequencing reads *τ*, the sequencing output *Y*_*gi*_. for all *i* were obtained as a multinomial with proportions vector 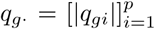. We set the control proportions from various experimental datasets (specifically, mouse, lung and the diarrheal microbiomes). With this setup, we can vary *ƒ*, and the two parameters of the fold change distribution, and ask, how various normalization and testing procedures compare in terms of their performance. For bulk RNAseq data, as illustrated in supplementary figure 1, we simulated 20M reads per sample.

For comparison of Wrench scales with other normalization approaches, we altered the above procedure slightly to allow for variations in internal abundances of features in observations arising from a group *g*. We used 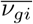 ( where the bar indicates this value will now assume the role of an average) generated above as a prior fold change for observation-wise fold change generation. That is, for all samples *j* ∈ 1 … *n*_*g*_ for all *g*, where *n*_*g*_ represents the number of samples in group *g*, for all *i* (including the truly null features), sample *v*_*gji*_ from 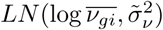 for a small value of 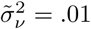. This induces sample specific variations in the proportions within groups. Notice that this makes the problem harder and more realistic, as feature marginal count distributions now arise from a mixture of distributions. Based on empirically observed MA plots for our metagenomic datasets, we set the mean and standard deviation of prior log-fold change distribution to 0 and 3 respectively. For generating 16S metagenomic-like datasets, logged sample depths were sampled from a log-normal distribution with logged-standard deviation of .25 and logged-means corresponding to log(4*K*), log(10*K*) and log(100*K*) reads. These parameters were chosen based on comparisons with MA plots, the sparsity levels and total sample depths observed in current experimental datasets.

In both versions of simulations, the total induced abundance change relative to that of the control is 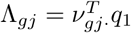, where *v*_*gj*_. is the vector of fold changes for sample *j*, and qi is the average vector of control proportions. We apply the term *compositional correction factor* for 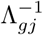 and the term *normalization factor* for a sample as the product of its compositional correction factor with something that is proportional to that of its sample depth. Thus, all technical artifacts like total abundance changes, but sample depth, are incorporated into the definition of compositional factors.

### Performance comparisons

For simulations, we used edgeR as the workhorse fitting toolkit. The compositional scale factors provided by all normalization methods were provided to edgeR as offset factors. We define detectable differential abundance in our simulated count data as follows. For each simulation, as we know the true compositional factors, we input them as normalization factors in edgeR, and the detectable differences in abundances are recorded. All the performance metrics are then defined based on this ground truth. Because we are interested in fold changes and their directions, the performance metrics we report are redefined as follows: Sensitivity as the ratio of the number of detectable true-positives with true sign over the total number of positives, False discovery as the ratio of the number of detectable true positives with false sign and false positives, over the total number of significant calls made.

The offset-covariate analysis followed the procedure in [25]. For resampling analysis, samples from each experimental group (with atleast 15 samples) were split in half randomly to construct two artificial groups. Normalization factors from each method were then used to perform differential abundance analysis, and the total number of differentially abundant calls were recorded. The procedure was repeated for ten iterations for each group, and the results were averaged across 41 experimental groups. Those samples for which Scran fails to reconstruct normalization scales were discarded from differential abundance analyses to avoid any power differences while testing. The normalization scales however, were obtained with all data for each method.

Fisher exact tests were used to perform functional enrichment analyses for positively associated OTUs. A Genus level functional enrichment analysis was first performed by aggregating annotations from the FAPROTAX1.1 database [43] at the Genus level. A more specific OTU level functional enrichment analysis was devised as follows. Because the Tara Oceans Kegg module (KM) abundance data (downloaded from http://ocean-microbiome.embl.de/data/TARA243.KO-module.profile.release.gz) and the 16S reconstructions are obtained from the same input DNA through whole metagenome shotgun, the same compositional factors apply to both datatypes. Each normalization approach’s compositional factors for 16S data was used to rescale the KM relative abundance data. This normalized KM data was used to annotate each OTU by (normalized) KMs that Spearman correlate at a value of atleast .75.

### Software availability

Wrench is available from GitLab as an R package at the URL: https://gitlab.umiacs.umd.edu/smuthiah/Wrench.

## Competing interests

The authors declare that they have no competing interests.

## Author’s contributions

Conceived and designed study: MSK, HCB.

Contributed analytical tools/reagents: MSK, EVS, KO, HCB.

Wrote the paper: MSK, SH, HCB.

Participated in discussions: All authors.

## Acknowledgements

MSK thanks Mihai Pop, Tom Goldstein, Joyce Hsiao and Mathieu Almeida for useful discussions, Joseph Paulson for making the metagenomic count data used in this paper available through R/Bioconductor, the Tara Oceans and the HMP teams for making their processed count data easily available. This work was partially supported by NSF grant 1564785 to SH, by NIH R01 grants GM083084, RR021967/GM103552 and K99HG009007 to SCH, and by NIH R01 grants GM114267 and HG005220 to HCB and MSK.

## Additional Files

### Additional file 1 — Supplementary Note

Presents further discussions on compositional bias, and supplementary results in context.

### Additional file 2 — Enrichment Analysis Results

The results of enrichment analyses based on faprotax annotations and Kegg modules procedure described in the Methods section is presented. Names in the sheets and their descriptions are as follows: KM.POS.SIG.MES and KM.POS.SIG.DCM show the Kegg module based enrichment analyses for positively associated features in MES and DCM layers respectively. FAPRO.POS.SIG.MES and FAPRO.POS.SIG.DCM show the results of faprotax annotations based enrichment analyses for positively associated features in MES and DCM layers respectively.

## Footnotes

[1] the idea being that in the limit Λ_*g*_ → ∞, feature-wise ratios that reflect 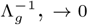

[2] the random variable assumes a value of zero with probability *π* and a positive value based on its specific log-normal distribution with probability (1 − *π*)

